# Neurometabolic correlates of accelerated aging and neurocognitive late effects in long-term survivors of pediatric hodgkin lymphoma and acute lymphoblastic leukemia

**DOI:** 10.64898/2026.02.17.706361

**Authors:** Kyla Gibney, Areeb Khan, Sabah Nisar, Kasturee Chakraborty, Ritambhar Burman, Pat Hanby, Stephanie Guthrie, Brian Potter, Melissa M. Hudson, Kirsten K. Ness, Tara Brinkman, Belinda Mandrell, Cai Li, Kevin Krull, Puneet Bagga

## Abstract

Adult survivors of pediatric cancers are at elevated risk for neurocognitive late effects, but how these effects relate to metabolic perturbations in the brain remains unclear. To address this knowledge gap, the present study explored associations between neurometabolite levels and neurocognitive function in adult survivors of Hodgkin lymphoma (HL) and acute lymphocytic leukemia (ALL). Data were collected from a single-center observational study conducted at St. Jude Children’s Research Hospital (SJCRH) between October 2022 and November 2024. Adult survivors of <u>HL</u> <u>(*N*=11 [5 females]; ≥5 years post-diagnosis; mean [SD] current age 34 [9.5] years)</u> and <u>ALL (*N*=24 [16 females]; ≥5 years post-diagnosis; current age 40 [12.6] years)</u> and <u>community controls (*N*=35 [17 females]; current age 40 [11] years)</u> completed standardized neurocognitive tests of memory, attention, executive function, and processing speed. Participants also underwent proton magnetic resonance spectroscopy (^1^H MRS) to quantify neurometabolite levels in the left dorsolateral prefrontal cortex (dlPFC), left hippocampus, and left cerebellum. Analyses used regression models to examine differences in the slope of the relationship between neurometabolite and neurocognitive function or between neurometabolite and age. When comparing HL survivors vs controls, significant interactions were identified for group x age on the ratio of myo-inositol to N-Acetyl aspartic acid (mI/NAA; *p*=0.007) and group x Gamma-Aminobutyric Acid (GABA) on processing speed (*p*=0.04) in the left dlPFC. When comparing ALL survivors vs controls, significant interactions were identified for group x myo-inositol on verbal fluency in the left hippocampus (*p*=0.01) and group x GABA on cognitive flexibility in the left cerebellum (*p*=0.01). These preliminary findings suggest that neuroinflammation may be a mechanistic underpinning of age-associated neurocognitive impairment in pediatric cancer survivors.

## 1. INTRODUCTION

Hodgkin Lymphoma (HL) and acute lymphocytic leukemia (ALL) comprise nearly 50% of pediatric cancer cases in the United States^1,2^. Although the 5-year survival rate for pediatric cancers now exceeds 85%^3,4^, cancer treatments such as chemotherapy and radiation therapy (RT) can lead to adverse health outcomes later in life, such as cardiopulmonary and thyroid dysfunction^5–8^, neurocognitive impairment^9–12^, epigenetic age acceleration^13,14^, and chronic systemic inflammation^15,16^. Some of these late effects, such as chronic inflammation and neurocognitive impairment, are also part of the typical aging process; however, they may be more pronounced in cancer survivors, for whom the normative aging process exacerbates the late effects of cancer treatment. These treatment-related late effects pose a substantial burden to the over 100,000 survivors of pediatric HL and ALL in the US today^1–3,10^. This is particularly relevant for individuals diagnosed during childhood who have decades of survival post-diagnosis. To improve survivors’ quality of life, a deeper understanding of the mechanisms underlying cognitive dysfunction following pediatric cancer is needed.

We previously reported that cognitive deficits in pediatric cancer survivors are linked to observable brain pathology. In HL survivors, cerebrovascular injury and multifocal leukoencephalopathy are common sequelae, and these effects appear alongside cardiopulmonary dysfunction, particularly in those who received higher-dose thoracic RT^12,17,18^ Neuroimaging of pediatric ALL survivors has revealed both structural and functional alterations, such as reduced white matter integrity, altered default mode network connectivity, and altered blood oxygen level dependent (BOLD) responses in prefrontal, parietal, and hippocampal regions^9,18–20^. Furthermore, our group has found that survivors of pediatric cancer accrue age-associated physiological deficits at a rate that is much higher than the general population, and their DNA methylation signatures demonstrate signs of epigenetic EAA^14,21–23^.

Although it is known that adult survivors of both pediatric HL and ALL experience neurocognitive impairment as a result of their cancer treatment, there are likely distinct neurometabolic pathways underlying treatment-related neurocognitive decline between these two groups. In the case of aging or pathology, metabolic changes may occur before any observable structural alterations manifest in the brain^24–27^. Thus, by elucidating the dynamics of neurometabolism during aging, we may be able to identify the development of pathologies before they become visible on a functional or anatomical level.

The present study explores the association between neurocognitive late effects following cancer treatment with neurometabolism in survivors of pediatric HL and ALL compared to community controls using proton magnetic resonance spectroscopy (^1^H MRS), a non-invasive technique used to quantify neurometabolites. Multiple studies have demonstrated that neurometabolism and neurocognitive function change over the lifespan in general populations^10–13^. Given the advanced aging observed in pediatric cancer survivors, we hypothesized that they would exhibit a stronger association between age-related metabolites and their chronological age compared to community controls, and we expected that neurotransmitter-related metabolites, such as glutamate (Glu) and glutamine (Gln), would be associated with neurocognitive performance in survivors.

## 2. METHODS

### 2A. Study Participants

The data presented in the current study were collected as part of ancillary studies in the St. Jude Lifetime Cohort Study (SJLIFE). SJLIFE is a retrospective cohort study with prospective follow-up of patients with malignancies diagnosed and treated at St. Jude Children’s Research Hospital (SJCRH) between 1962–2012 who are ≥5 years post-completion of treatment^28^. The Institutional Review Board approved the study, and participants or their guardians provided written informed consent.

The present study includes adult survivors of pediatric HL and ALL who were treated with chest RT, cranial RT, and/or chemotherapy. Survivors were ≥18 years old at the time of their evaluation. Community controls without a history of cancer were recruited as part of a larger study to frequency-match survivors by age, sex, race, and body mass index. Participants meeting any of the following criteria were excluded from the study: a history of head injury; a genetic disorder associated with neurocognitive impairment; a birth complication associated with neurocognitive impairment; a history of congenital heart disease; or current pregnancy. See **Table 1** for demographic information, including age, sex, and patient population. Treatment exposures for the HL and ALL survivors are outlined in **Table 2**.

**Table 1:** Participant Demographics. Demographic information for ALL survivors, HL survivors, and control subjects, including sex, age at the time of assessment, age at the time of diagnosis, and how many years had elapsed since the HL and ALL survivors were diagnosed.

| **Table 1. Participant Demographics** |  |  |  |  |  |  |
| --- | --- | --- | --- | --- | --- | --- |
|  | **ALL survivors**<br>**(N=24)** | **HL survivors**<br>**(N=11)** | **p-val**<br>**(ALL vs HL)** | **Controls**<br>**(N=35)** | **p-val**<br>**(ALL vs controls)** | **p-val**<br>**(HL vs controls)** |
| **Age at assessment, years** |  |  |  |  |  |  |
| Mean ± SD | 40 ± 12.6 | 34 ± 9.5 | 0.1 | 40 ± 11.3 | 0.001 | 0.1 |
| **Age at diagnosis, years** |  |  |  |  |  |  |
| Mean ± SD | 7.6 ± 4.8 | 12.8 ± 4.9 | 0.01 | ----- | ----- | ----- |
| **Time since diagnosis, years** |  |  |  |  |  |  |
| Mean ± SD | 33 ± 11.1 | 21.7 ± 8 | 0.005 | ----- | ----- | ----- |
| **Sex, n (%)** |  |  |  |  |  |  |
| Female | 16 (66%) | 5 (46 %) | 0.2 | 17 (49%) | 0.2 | 0.9 |
| Male | 8 (33%) | 6 (54 %) |  | 18 (51%) |  |  |

**Table 2:** Treatment Exposures. Treatment exposures for ALL and HL survivors, including the cumulative dose of alkylating agents, anthracyclines, chest radiotherapy, and cranial radiotherapy.

| Table 2. Treatment Exposures |  |  |  |
| --- | --- | --- | --- |
|  | ALL survivors<br>(N=24) | HL survivors<br>(N=11) | p-val |
| **Chemotherapy, cumulative dose** |  |  |  |
| **Alkylating agents (mg/m<sup>2</sup>)** |  |  |  |
| Mean ± SD (n) | 7492 ± 4367 (n=20) | 5486 ± 6462 (n=9) | 0.4 |
| **Anthracyclines (mg/m<sup>2</sup>)** |  |  |  |
| Mean ± SD (n) | 91 ± 59 (n=19) | 158 ± 18 (n=9) | 0.0001 |
| **Chest Radiotherapy dose (cGy)** |  |  |  |
| Mean ± SD (n) | 268 ± 784 (n=10) | 2071 ± 915 (n=7) | 0.001 |
| **Cranial Radiotherapy dose (cGy)** |  |  |  |
| Mean ± SD (n) | 2000 ± 326 (n=10) | 20 ± 0 (n=8) | <<0.0001 |

### 2B. MR Scanning

MRI scanning was performed on a 3T MRI scanner (Siemens Medical Systems, Erlangen, Germany). Acquisition of ^1^H MRS data was preceded by a T1-weighted MPRAGE scan (TR/TE/ 6.9/3.2 ms; FA 8°) with 1 mm^3^ isotropic resolution for voxel positioning and tissue segmentation. PRESS ^29^ localization was used to acquire ^1^H MRS with voxels positioned in the left dorsolateral prefrontal cortex (dlPFC), left hippocampus, and left cerebellum (**Figure 1**). We focused on the left dlPFC as our primary region of interest due to its prominent role in top-down attention and executive function, which are commonly impacted by aging^30–32^. We also included the left hippocampus due to its prominent role in learning, memory, and language processing ^33–36^. Meanwhile, the cerebellum was included due to its role in motor coordination and its prominent connectivity with top-down executive control networks in the dlPFC^37^.

**Figure 1:**
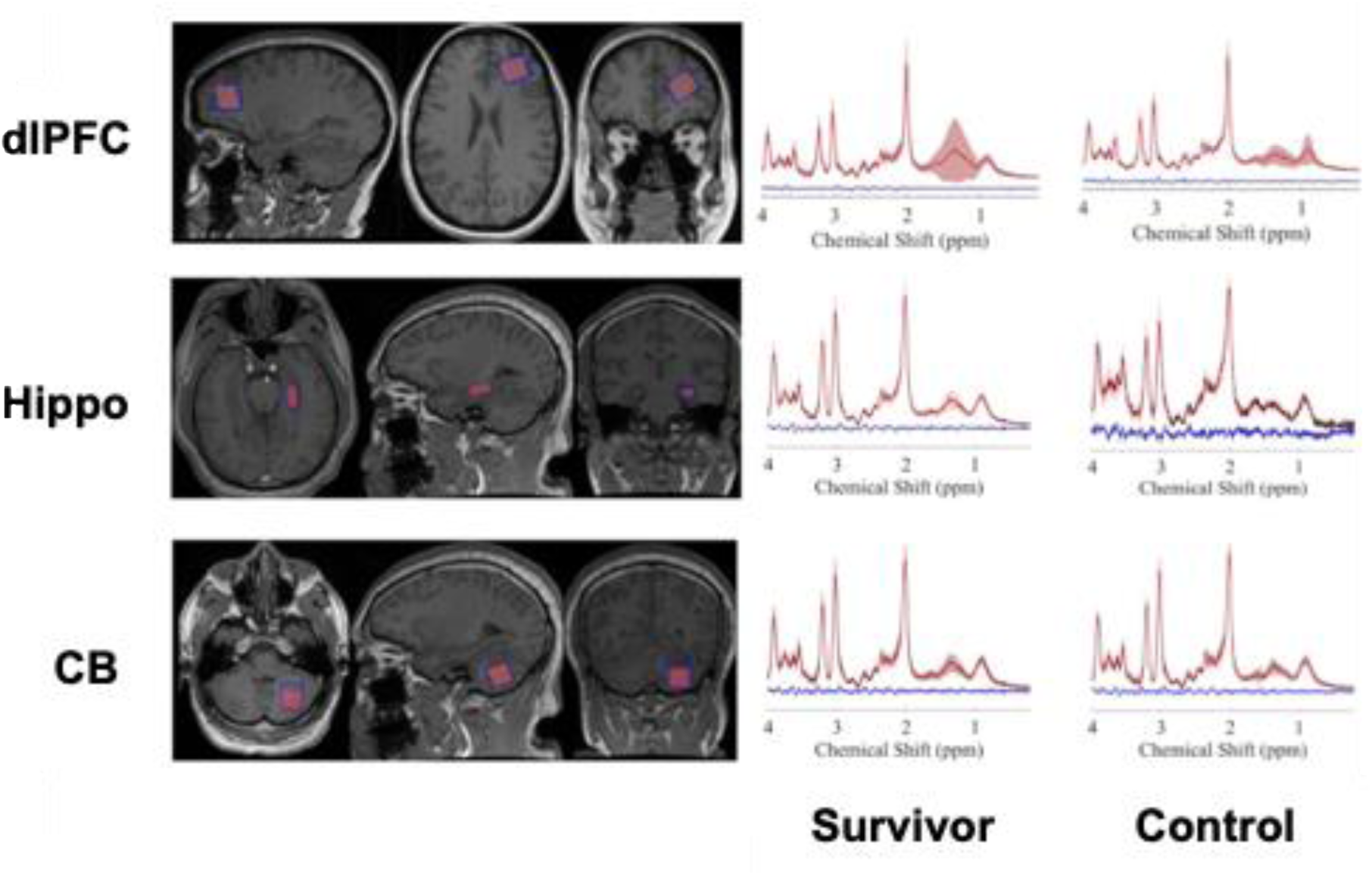
Example MRS voxel placement (red) in the left dlPFC, left hippocampus, and left cerebellum on an anatomical image (above left). Average raw spectra (black) obtained with ^1^H MRS in survivors (left) and controls (right) for the left dlPFC, left hippocampus, and left cerebellum (above right). Average fitted data (red) and average residuals (blue) are also shown. The shaded areas represent ± SD. ppm: parts per million; dlPFC: dorsolateral prefrontal cortex; Hippo: hippocampus, CB: cerebellum.

Raw ^1^H MRS data files were transferred offline for fully automated postprocessing using an in-house software tool called High Throughput LCModel (HT-LCModel) ^38^. HT-LCModel provides a high-throughput, GUI-driven pipeline for analyzing ^1^H-MRS data in Windows environments using the Linux-based LCModel software^39^. LCModel provides objective measures for the signal-to-noise ratio (SNR) and the spectral line width (full width at half maximum) for objective quality assessment. Spectra were fitted between 0.2 and 4.0 ppm, and spectra with low SNR were discarded automatically using HT-LCModel. Levels of N-acetyl aspartic acid (NAA), choline-containing compounds (Cho), total creatine (tCr), glutamate (Glu), glutamate + glutamine (Glx), gamma-aminobutyric acid (GABA), myo-inositol (mI), and glutathione (GSH) were determined. Metabolite ratios with a Cramér-Rao Lower Bound (CRLB)≤30 were included in the analysis. Metabolites are expressed either as a ratio relative to total creatine (/tCr) or to another metabolite (for example, mI/NAA). Henceforth, the metabolite ratio to creatine will be represented as the metabolite itself.

### 2C. Neurocognitive function

The participants underwent standardized neurocognitive testing on the same day that their MRS data was acquired. Assessments included tests of memory (California Verbal Learning Test [CVLT]), executive function (Digit Span Backwards, Verbal Fluency Test, and Part B of the Trail Making Test [TMT B]), and processing speed (Digit Symbol Coding Task) because these domains are commonly impacted during the aging process and by cancer survivorship^40–44^.

### 2C. Statistical Analysis

Statistical analyses were performed in R version 4.1^45^. The Mann-Whitney test was used to examine the difference in metabolite levels between control subjects and cancer survivors (ALL or HL). We used t-tests to determine differences in age (**Table 1**), treatment exposures (**Table 2**), neurocognitive performance (**Table 3**), and neurometabolite levels (**Figure 2**) between groups. We used Chi-squared tests to determine differences in sex (**Table 1**) between groups.

**Figure 2:**
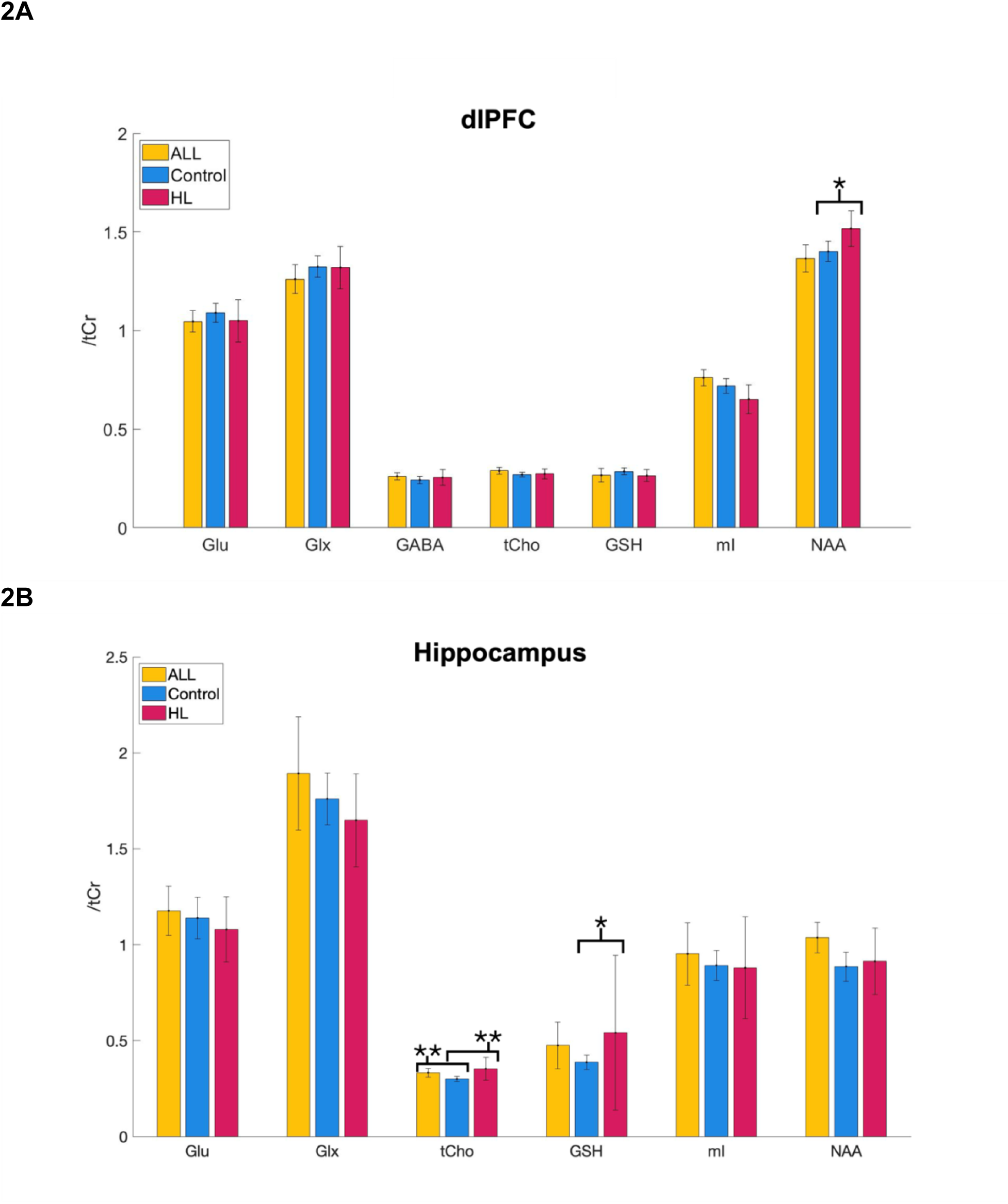
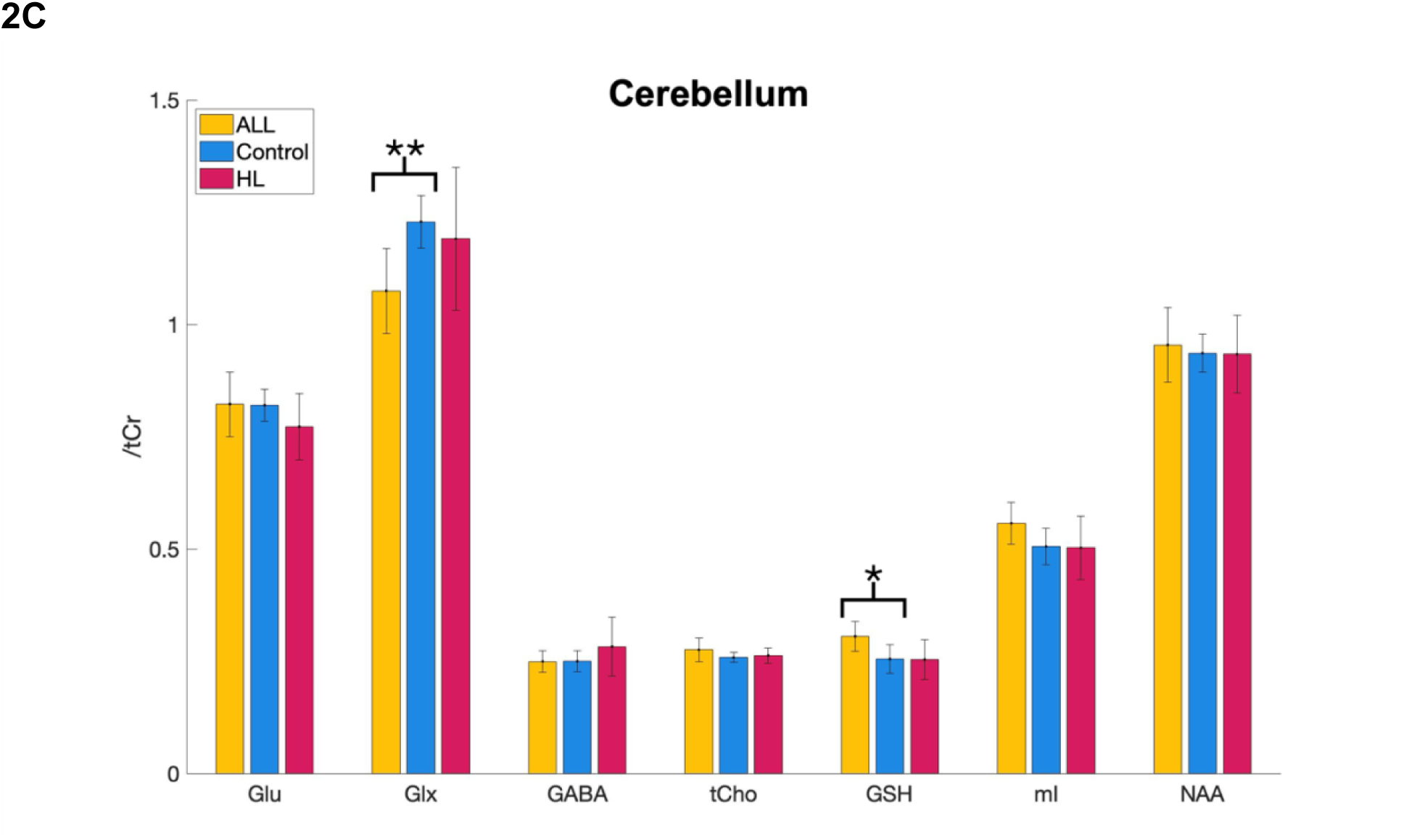
A comparison of neurometabolite levels between ALL survivors, HL survivors, and control subjects in the dlPFC (**A**), hippocampus (**B**), and cerebellum (**C**). dlPFC: dorsolateral prefrontal cortex, ALL: acute lymphocytic leukemia, HL: Hodgkin lymphoma, Glu: glutamate, Glx: glutamate + glutamine, GABA: gamma-amino butyric acid, tCho: choline-containing compounds, GSH: glutathione, mI: myo-inositol, NAA: N-acetyl aspartic acid, tCr: total creatine. Error bars indicate 95% confidence intervals. \**p*< 0.05, \*\**p*<0.01

**Table 3:** Neurocognitive Performance. Performance on the Trail Making, Verbal Fluency, Digit Symbol Coding, Digit Backwards, and California Verbal Learning tests among ALL survivors, HL survivors, and control subjects. ^a^measure of cognitive flexibility ^b^measure of verbal fluency ^c^measure of processing speed ^d^measure of working memory ^e^measure of verbal memory

| **Table 3. Neurocognitive Performance** |  |  |  |  |  |
| --- | --- | --- | --- | --- | --- |
| Z-score: Mean ± SD | **ALL survivors**<br>(N=24) | **HL survivors**<br>(N=11) | **Controls**<br>(N=35) | **p**<br>(ALL vs controls) | **p**<br>(HL vs controls) |
| **Trail Making Test, Part B<sup>a</sup>** | 0.19 ± 1.13 | -0.53 ± 1.61 | 0.35 ± 0.93 | 0.6 | 0.03 |
| **Verbal Fluency<sup>b</sup>** | 0.18 ± 1.17 | -0.3 ± 1.04 | 0.039 ± 0.99 | 0.6 | 0.3 |
| **Digit Symbol Coding<sup>c</sup>** | 0.14 ± 0.96 | -0.18 ± 1.43 | 0.49 ± 0.87 | 0.2 | 0.07 |
| **Digit Backwards<sup>d</sup>** | -0.15 ± 0.91 | -0.48 ± 0.91 | -0.088 ± 0.8 | 0.8 | 0.2 |
| **CVLT, Total Score<sup>e</sup>** | -0.51 ± 1.07 | -0.44 ± 1.35 | 0.36 ± 0.95 | 0.6 | 0.03 |

We used linear regression to examine the effect of group (survivors vs controls) and age, as well as interactions, adjusting for sex, on each neurometabolite in the left dlPFC. Similar regressions were used to quantify the effect of group and neurometabolite on neurocognitive function. A significant interaction effect from these tests was interpreted as a difference in the slope of the relationship between metabolite(s) and variable (age or neurocognitive function) between groups. Multiple comparisons correction was conducted using the Bonferroni method.

## 3. RESULTS

The sample included 11 adult survivors of pediatric HL (mean [SD] age at evaluation 34 [9.5] years; 5 females [46%]), 24 adult survivors of pediatric ALL (40 [12.6] years; 16 females [66%]), and 35 community controls (40 [11.3] years; 17 females [49%]). ALL survivors were diagnosed at a significantly younger age than HL survivors (*p*=0.01), and, accordingly, significantly more time had passed since diagnosis for ALL survivors compared to HL survivors (*p*=0.005). Demographic information for these participant samples is outlined in **Table 1**.

Among our sample of 24 ALL survivors, 20 were treated with alkylating agents (mean [SD] cumulative dose: 7492 [4367] mg/m^2^), 19 were treated with anthracyclines (91 [59] mg/m^2^), and 10 were treated with chest RT (mean [SD] max dose: 268 [784] cGy) and cranial RT (2000 [326] cGy). Among our sample of 11 HL survivors, 9 were treated with alkylating agents (5486 [6462] mg/m^2^) and anthracyclines (158 [18] mg/m^2^), 7 were treated with chest RT (2071 [915] cGy), and 8 (20 [0] cGy) were treated with cranial RT. HL survivors were exposed to significantly higher doses of anthracyclines (*p*=0.0001) and chest radiotherapy (*p*=0.001) compared to ALL survivors; meanwhile, ALL survivors were exposed to higher doses of cranial radiation compared to HL survivors (*p*<<0.0001; **Table 2**).

HL survivors showed significantly poorer performance on cognitive flexibility (part B of the Trail Making Task, *p*=0.03) and the verbal memory (CVLT, *p*=0.03) compared to control subjects (**Table 3**). In the left dlPFC, NAA was significantly higher among HL survivors than control subjects (*p*=0.02; **Figure 2A**). In the left hippocampus, tCho (*p*=0.005) and GSH (*p*=0.03) were significantly higher for HL survivors than control subjects, and tCho was significantly higher among ALL survivors than controls (*p*=0.007; **Figure 2B**). In the left cerebellum, GSH was significantly higher among ALL survivors than controls (*p*=0.03) and Glx was significantly higher among controls than ALL survivors (*p*=0.004; **Figure 2C**).

When comparing HL vs control subjects, we found a significant interaction effect for Group x Age on mI/NAA (*p*=0.007) in the left dlPFC (**Figure 3A**). However, the interaction effects for Group x Age on mI (*p*=0.1) or NAA (*p*=0.1) were not significant (**Figure 3B-C**). We also found a significant interaction effect for Group (HL vs controls) x GABA on processing speed (Digit Symbol Coding, **Figure 4A**, *p*=0.04) in the left dlPFC.

**Figure 3:**
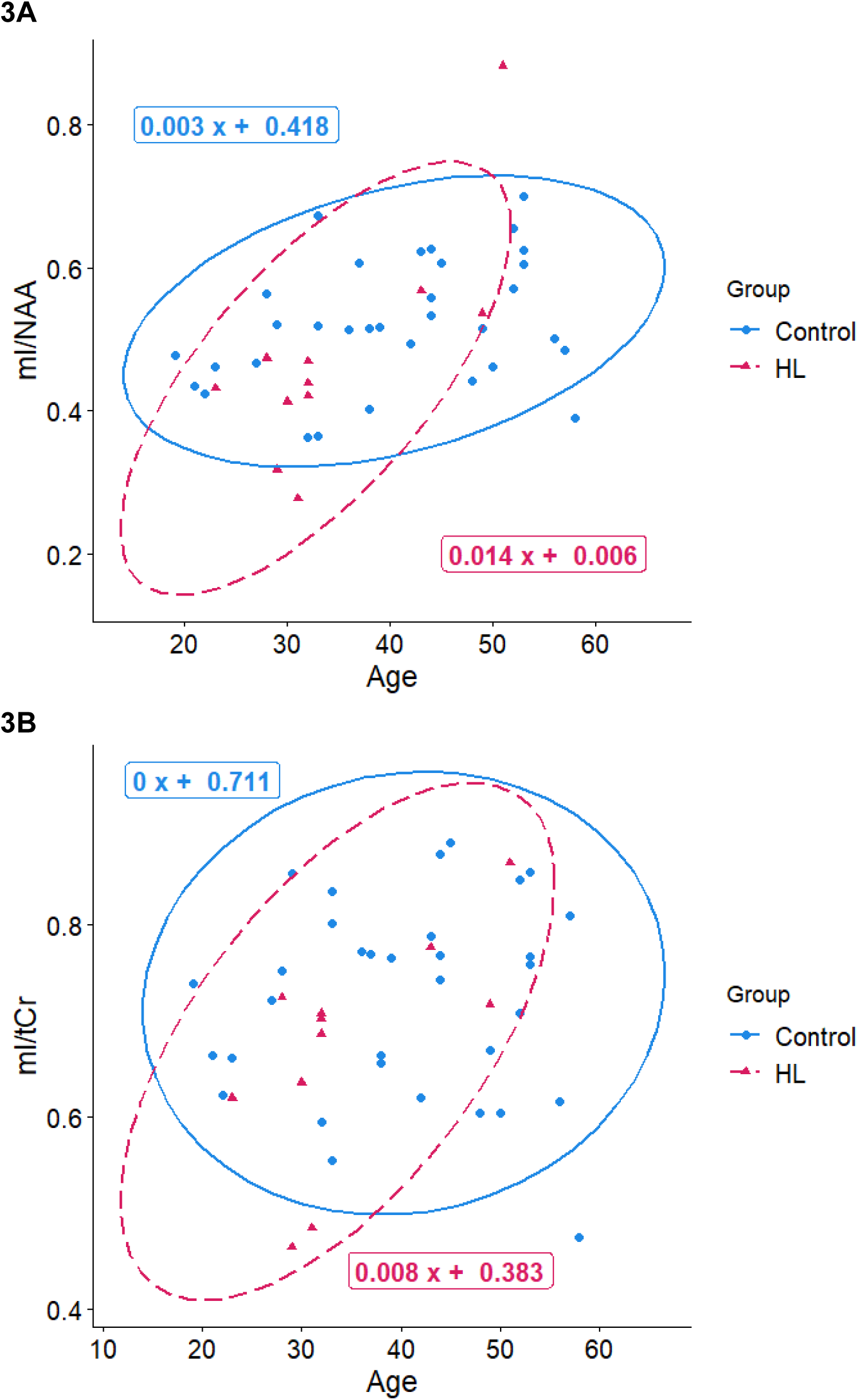
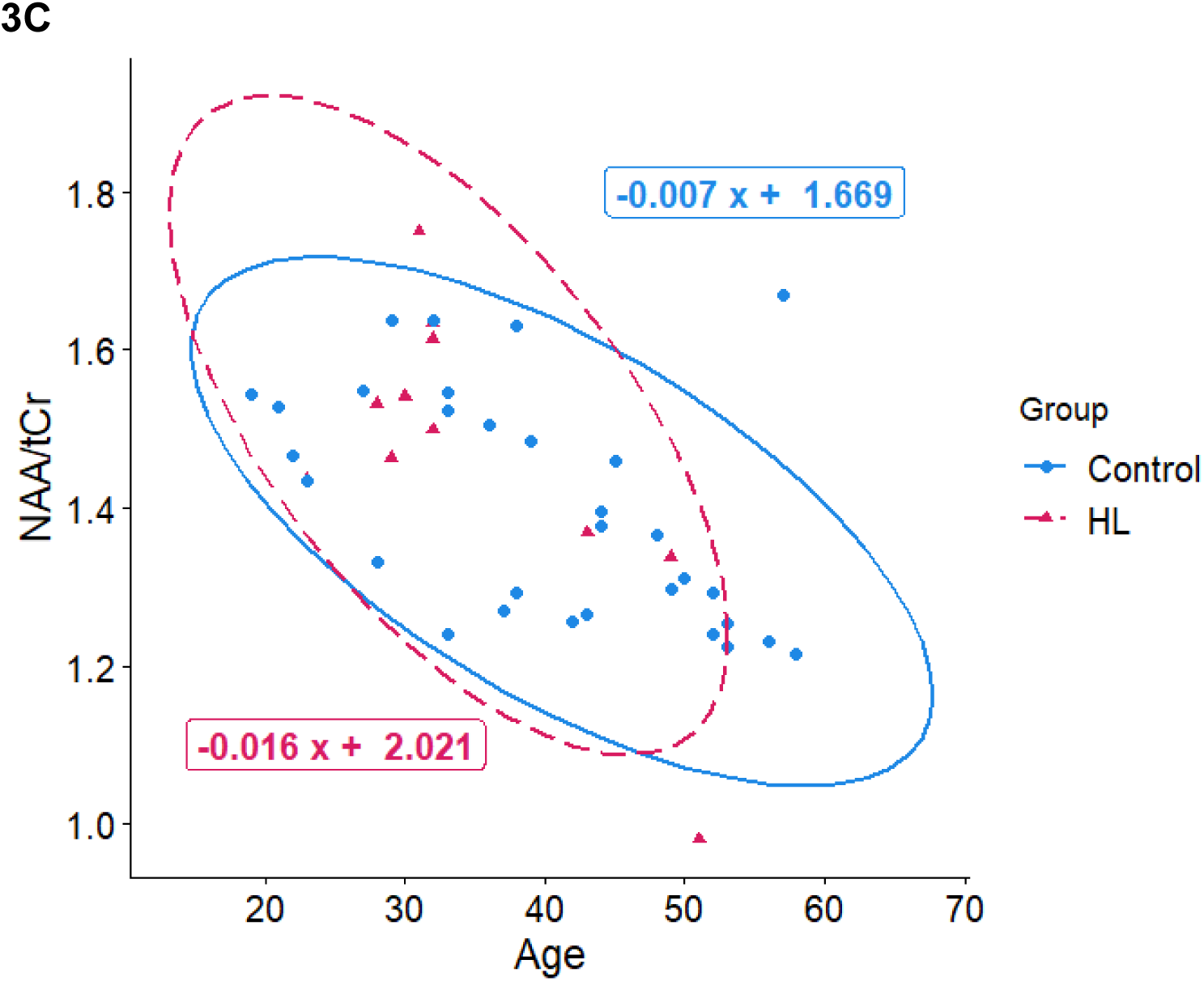
The association between age and mI/NAA (**A**), mI/tCr (**B**), and NAA/tCr (**C**) in the dlPFC for HL survivors (red triangles) and control subjects (blue dots). dlPFC: dorsolateral prefrontal cortex, mI: myo-inositol, NAA: N-acetyl aspartic acid, tCr: total creatine, HL: Hodgkin’s lymphoma.

**Figure 4:**
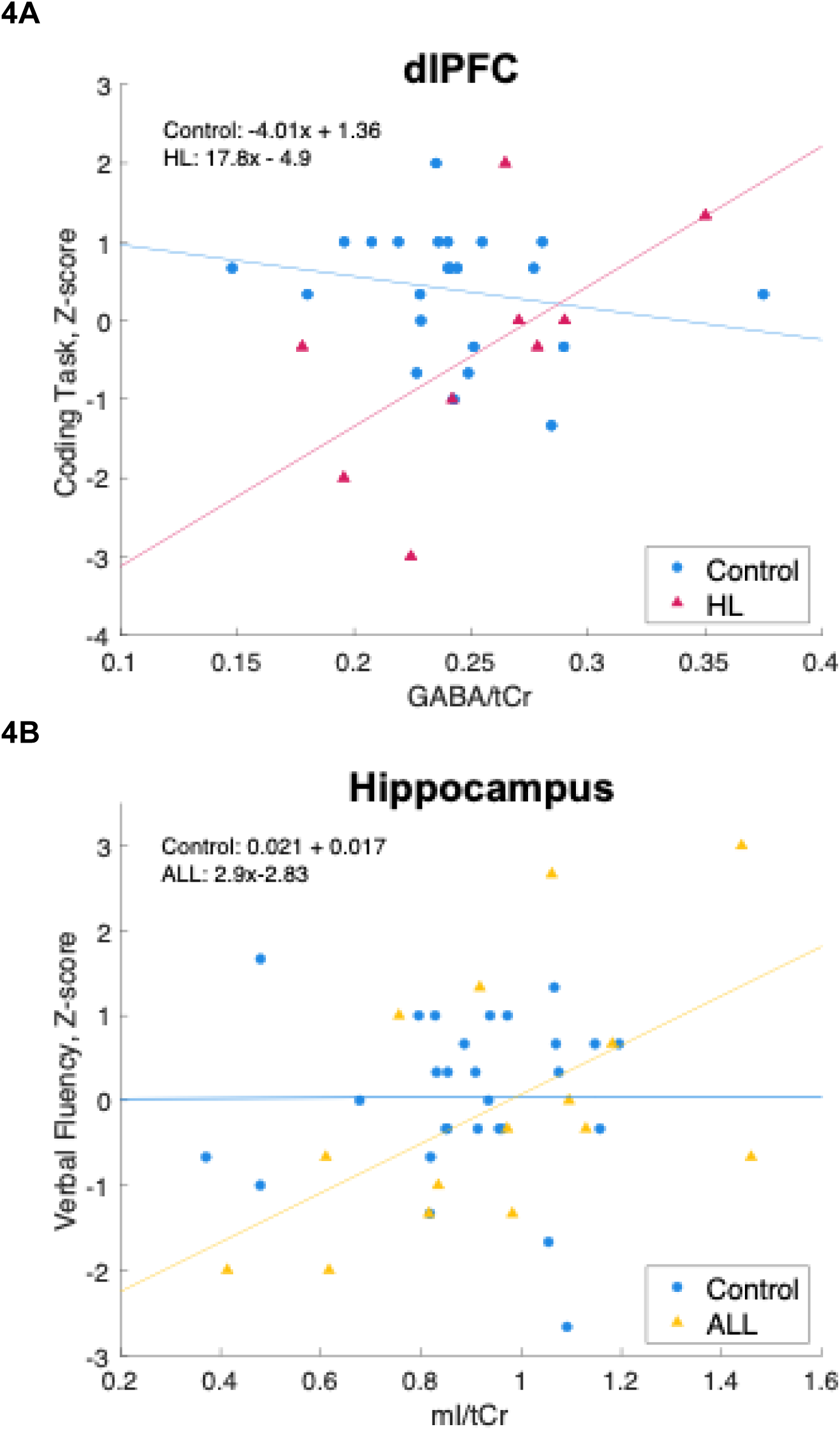
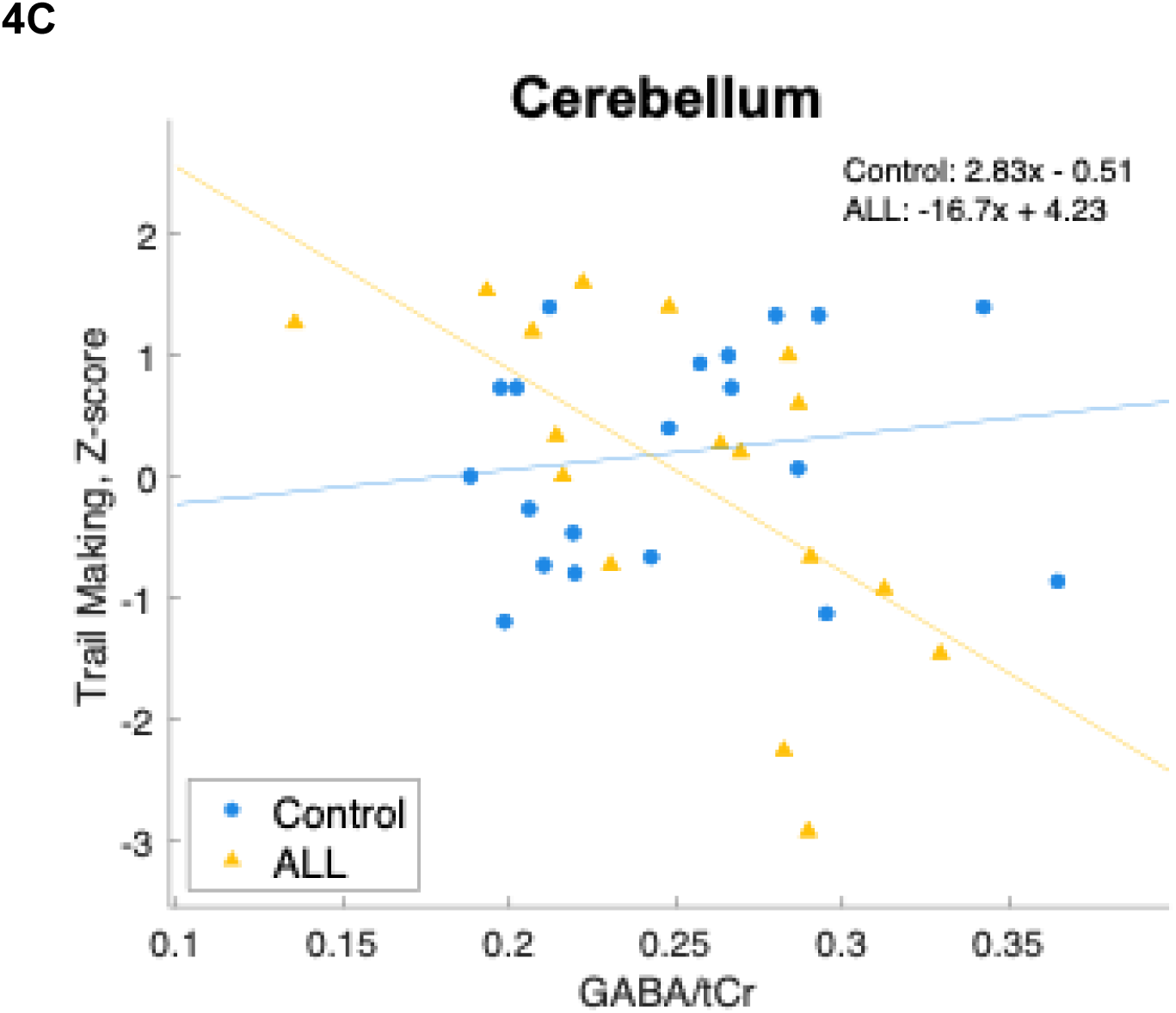
The association between performance on the Digit Symbol Coding Task and GABA/tCr (**A**), the Verbal Fluency Task and mI/tCr (**B**), and the Trail Making Task and GABA/tCr (**C**) for HL survivors (red triangles), ALL survivors (yellow triangles), and control subjects (blue dots). dlPFC: dorsolateral prefrontal cortex; GABA: gamma amino butyric acid; tCr: total creatine; HL: Hodgkin Lymphoma; mI: myo-inositol; ALL: acute lymphocytic leukemia; TMT B: Trail Making Task, Part B.

We identified age-related changes in the mI/NAA ratio in the left dlPFC of HL survivors that were not present in control subjects. For HL survivors, prefrontal cortical levels of mI increased with age (**Figure 3B**; *β*=0.083, *p*=0.03), whereas control subjects showed no such association (**Figure 3B**; *β≍*0, *p*=0.9). NAA also decreased with age among HL survivors (**Figure 3C**; *β*= −0.016, *p*=0.01) at a faster rate than among control subjects (**Figure 3C**; *β* = 0.007, *p*=0.001). Taken together, this creates a strong interaction effect for the ratio of mI/NAA between groups with age (**Figure 3A**).

When comparing ALL vs control subjects, we found a significant interaction effect of <u>Group x mI on verbal fluency in the left Hippocampus (</u>**Figure 4B**<u>; *p*=0.01)</u> and a significant interaction effect of Group x GABA on cognitive flexibility (Trail Making Task, Part B) in the left cerebellum (**Figure 4C**<u>; *p*=0.01</u>). All the aforementioned effects were significant after controlling for age and sex and correcting for multiple comparisons.

## 4. DISCUSSION

The present study explored the association(s) between age, neurocognitive function, and neurometabolism among a cohort of ALL survivors, HL survivors, and control subjects. Broadly, we found age-associated changes in the mI/NAA ratio that were different for survivors vs control subjects. We also found that the association between mI and GABA on verbal fluency and cognitive flexibility, respectively, differed between survivors and control subjects.

It is established that NAA decreases with age in the general population: NAA is a neuronal marker, and, with increasing age, there is often a loss of neurons that is part of the normal aging process^46,47^. However, the finding that NAA decreases at a faster rate for HL survivors than control subjects may indicate an elevated rate of brain aging among survivors. This result is consistent with previous findings from our group demonstrating accelerated brain aging in adult survivors of pediatric cancer ^11^. Our group has identified many DNA methylation signatures associated with epigenetic age acceleration (EAA) in T cells of cancer survivors, especially among genes regulating inflammatory processes^13^. Furthermore, these EAA alterations show strong correspondence to the many physiological deficits that survivors accumulate as they age^23^.

We found that mI increases with age in HL survivors (*β*=0.083, *p*=0.03), but there was no correlation between mI and aging in our control subjects (*β≍*0, *p*=0.9; **Figure 3B**). Because mI is a glial marker, and microglia are the primary immune cells of the brain, it is possible that elevated mI could indicate a heightened neuroinflammatory response. Taken in conjunction with the finding that many cancer survivors demonstrate chronic systemic inflammation^15,16^, it is feasible that these survivors experience ongoing central nervous system (CNS) inflammation that accelerates aging and age-associated cognitive decline.

The present study also found that performance on a verbal fluency task increases with hippocampal mI among ALL survivors (*β*=2.9, *p*=0.03). Meanwhile, we found no association between hippocampal mI and verbal fluency among control subjects (β=0.021, *p*=1). This created a strong interaction effect of Group x mI on verbal fluency for ALL survivors vs controls (**Figure 4B**; *p*=0.01).

The hippocampus plays a prominent role in synaptic plasticity and episodic memory ^35,48,49^. During language processing, the hippocampus may integrate inputs from other sources, such as multisensory signals from audiovisual areas or representational/semantic information from prefrontal areas ^35,36^. This often enables flexibility and fluency in speech processing ^33^. Thus, our finding that verbal fluency, which engages both memory and speech-related mechanisms^44^, shows an association with hippocampal neurometabolism, is consistent with many other groups’ findings regarding memory and speech in hippocampus. The association with hippocampal mI could indicate that ALL survivors rely on hippocampal glial activity to a higher degree than the general population when processing speech signals.

Increased levels of brain mI are likely to be multifactorial and may be associated with several underlying processes, such as neuroinflammation, glial activation, or the aging process itself^50^. Our group has found that elevated levels of systemic inflammatory markers were associated with worse neurocognitive performance^16,51^ among pediatric cancer survivors. Current results suggest this peripheral inflammation may cross the blood-brain barrier and lead to inflammation of the CNS. It is not fully understood how chest RT may contribute to this CNS inflammation, i.e., if it is a secondary effect of cardiopulmonary morbidity caused by chest RT or if it represents a sustained inflammatory response in the aftermath of treatment.

Although mI is generally considered a glial marker, it does not necessarily distinguish between different types of glial cells^52^. Thus, these changes could be due to microglial activation, reflecting an inflammatory mechanism^53^, or they could indicate astrocytic activity, perhaps reflecting the clearing of cellular wastes or glutamine cycling^54,55^. This further emphasizes the need for more direct measures of neuroinflammation, which may delineate whether these alterations in mI are indeed related to astroglial activity or if they are related to a microglial response.

In addition to glial markers, we also found significant associations with GABA. Prefrontal cortical GABA was positively associated with processing speed in HL survivors (*β*=17.8, *p*=0.08) but not control subjects (*β*=-4.01, *p*=0.3; **Figure 4A**). Because GABA is an inhibitory neurotransmitter, this may indicate that survivors have an enhanced need for top-down inhibitory control mechanisms compared to the general population when completing cognitively demanding tasks. In light of the difficulties many HL survivors experience with top-down executive control^51,56^, these perturbations in prefrontal GABA further corroborate the idea that modes of higher-order cognitive processing, such as response inhibition, may be impaired in HL survivors.

Meanwhile, we also found that GABA in cerebellum was associated with cognitive flexibility (TMT B) for ALL survivors (*β*=-16.7, *p*=0.009) but not control subjects (*β*=2.83, *p*=0.5; **Figure 4C**). The cerebellum is implicated in motor processing and is also highly connected with prefrontal areas^37^.Thus, this finding may indicate a need for relay between top-down control mechanisms and a cerebellar motor response in order to engage cognitive flexibility mechanisms and update ones motors responses with correct goal-oriented processing. Perhaps the finding that GABA, the primary inhibitory neurotransmitter, is associated with this task probing cognitive flexibility reflects a need for appropriate activation/inhibition balance between prefrontal and cerebellar networks that is impaired in ALL survivors. It is known that ALL survivors commonly have underdeveloped prefrontal areas ^9,57,58^, which may prevent them from being able to successfully relay that top-down information between prefrontal cortex and cerebellum.

In our conceptual model of aging, late effects, and neurocognitive impairment among HL and ALL survivors (**Figure 5**), cancer treatments such as chemotherapy and RT cause cellular injury, leading to EAA and downstream alterations in T cells. This in turn leads to chronic systemic inflammation, which may cross the blood-brain barrier, causing inflammation of the CNS, which is reflected in elevated levels of brain mI shown in **Figure 3A-B** This contributes to accelerated brain aging, which is reflected in the decreased levels of NAA among survivors shown in **Figures 3A and C**. This accelerated brain aging also manifests in poorer performance on average on neurocognitive tests of attention, memory, executive function, and processing speed. Perhaps HL survivors must engage inhibitory control systems to a greater level to compensate for neurocognitive difficulties, which is reflected in increased prefrontal GABA among those with better task performance on the digit-symbol coding task (**see Figure 4A**).

**Figure 5:**
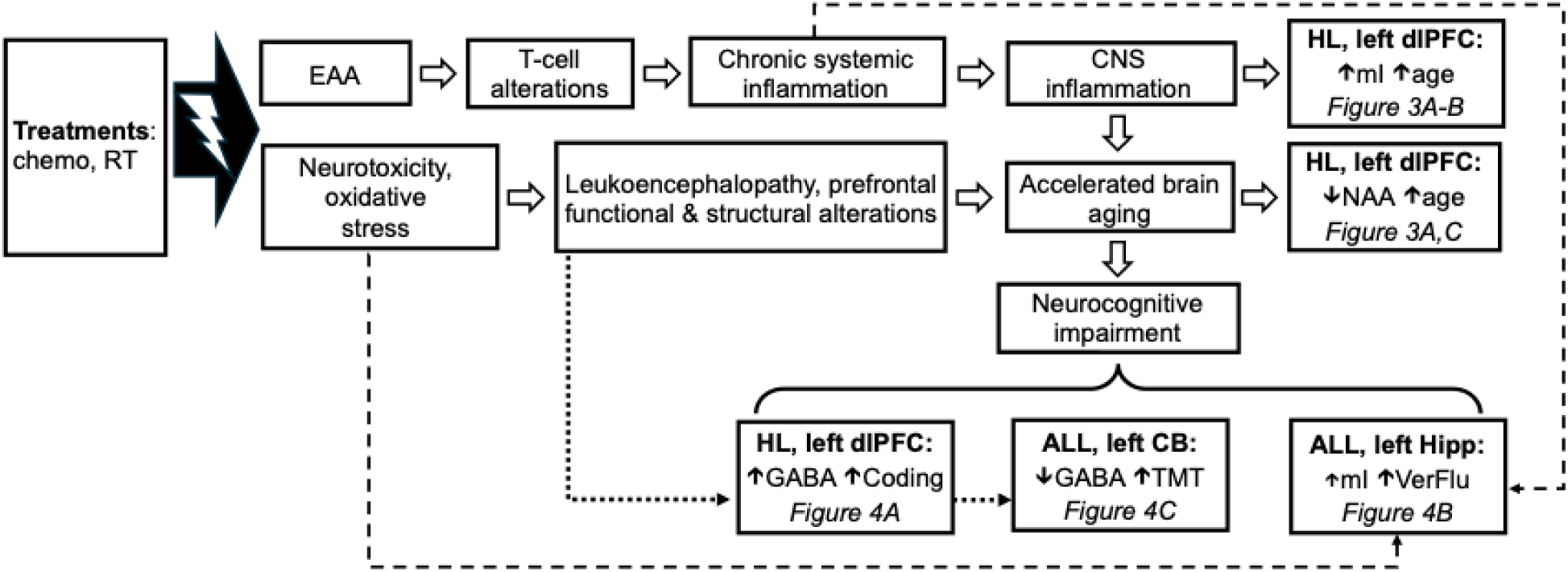
Conceptual model outlining the potential associations between cancer treatment, systemic inflammation, neuroinflammation, aging, and neurocognitive impairment in long-term survivors of pediatric cancer. Dashed lines indicate potential links between known molecular/inflammatory late effects, whereas dotted lines indicate potential links between known functional/anatomical late effects, that may provide a mechanistic explanation for our findings. RT: radiation therapy; EAA: epigenetic age acceleration; CNS: central nervous system; mI: myo-inositol; NAA: n-acetyl-aspartic acid; GABA: gamma-aminobutyric acid; TMT B: Trail-Making Test, Part B

In another stream of treatment-related late effects, chemotherapy and RT are neurotoxic and may cause oxidative stress. This in turn can lead to structural and functional alterations in brain networks necessary for normative cognitive function, especially prefrontal areas, which are highly connected to cerebellum. These sequelae also likely contribute to the accelerated brain aging and neurocognitive impairments seen in survivors of pediatric cancer. Namely, these structural and functional alterations may be reflected in the findings shown in **Figures 4A and 4C**, in which neurocognitive performance in survivors is associated with neurometabolite alterations in dlPFC and cerebellum, which are highly connected and implicated in top-down cognitive control.

The finding that ALL survivors with higher levels of hippocampal mI had higher Verbal Fluency performance (**Figure 4B**) suggests that ALL survivors may rely on a glial mechanism in order to effectively engage verbal fluency and memory networks in the hippocampus. It is possible that another mechanism in addition to neuroinflammation is responsible for the accelerated brain aging we observe in cancer survivorship, such as glutamine cycling in astrocytes.

Furthermore, the finding that ALL survivors with lower levels of cerebellar GABA had better TMT B performance (**Figure 4C**) could indicate the need for cognitive flexibility when completing this task and also underscores the prominent role of the cerebellum in not only motor function but also cognitive processing, due to its connectivity to prefrontal areas involved in top-down control. This difference also may be a downstream manifestation of the structural differences found in survivors of pediatric ALL, who often demonstrate leukoencephalopathy and underdeveloped prefrontal regions. Thus, as a consequence of pediatric cancer treatment, ALL survivors may have alterations in the relay between prefrontal and motor areas when conducting tasks with both motor and cognitive demands.

In our conceptual model of aging, late effects, and neurocognitive impairment among HL and ALL survivors (**Figure 5**), we hypothesize that cancer treatments such as RT and chemotherapy lead to EAA in survivors, causing immunomodulatory alterations that inflict cellular damage and lead to downstream inflammation. This systemic inflammation continues throughout the survivor’s life and may eventually cross the blood-brain barrier, where it may infiltrate the central nervous system and causes a neuroinflammatory response. This elevated neuroinflammatory response accelerates the normal aging process in the brain, leading to neuronal damage that may make it more difficult for survivors to engage in cognitively demanding tasks. Thus, in the context of survivorship, typical processes in the aging brain are exacerbated by the accumulated cellular insults of cancer treatment, eventually leading to neurocognitive impairment later in life.

## 5. LIMITATIONS

Our study is limited by small sample size, which hampers the ability to fully interpret the results and limits opportunities for multivariable modeling. Future studies with a larger participant sample are necessary to fully elucidate the relationship between age, neurocognitive impairment, and neurometabolite concentrations in HL and ALL survivors. Furthermore, the current findings do not offer insights into the distinct impact of cancer itself compared to that of cancer treatments on brain metabolite levels or other long-term effects on health. To elucidate whether the differences between groups reflect changes from their pre-illness state, longitudinal studies are necessary.

In this study GABA was detected using a non-edited PRESS MRS sequence. However, the accurate detection of GABA is feasible using editing i.e. MEGA-PRESS^59^. Finally, in order to precisely delineate the role of specific glial cell types in age-associated neurocognitive impairment during cancer survivorship, further studies employing more precise metrics of glial activity are necessary. For example, transmembrane protein (TSPO) is highly expressed on activated microglia, so positron emission tomography (PET) studies using TSPO-sensitive tracers may provide a more direct metric of neuroinflammation than ^1^H MRS alone^60–63^.

To fully interpret the present findings and their potential clinical implications, we must understand the underlying mechanisms driving these neurometabolic alterations. Further research spanning neuroimaging, clinical assessment, and biochemistry is warranted to delve deeper into the factors contributing to the alterations in neurometabolism among survivors.

## 6. CONCLUSIONS

Our study revealed that mI/NAA increases with age and GABA increases with processing speed performance in the dlPFC of HL survivors but not control subjects. We also found that mI decreases with cognitive flexibility among ALL survivors but increases with cognitive flexibility among control participants. These findings suggest that HL survivors may have an enhanced need for inhibitory control compared to the general population and that neuroinflammation may be a mechanistic mediator of age-related cognitive decline among HL survivors. Furthermore, our finding that hippocampal mI was associated with verbal fluency for ALL survivors further underscores a role of glial cells in neurocognitive function among pediatric cancer survivors.

Finally, our finding that cerebellar GABA was associated with cognitive flexibility for ALL survivors but not controls may indicate a need for appropriate relay between prefrontal and cerebellar regions in order to successfully engage cognitive flexibility. Thus, although the prefrontal cortex is highly connected to the cerebellum, in the case of ALL survivors, who are shown to have underdeveloped prefrontal areas, it is possible that the typical activation/inhibition balance may be dysregulated compared to control subjects.

In our conceptual model of aging, late effects, and neurocognitive impairment among HL and ALL survivors (**Figure 5**), we hypothesize that cancer treatments such as RT and chemotherapy lead to EAA in survivors, causing immunomodulatory alterations that inflict cellular damage and lead to downstream inflammation. This systemic inflammation continues throughout the survivor’s life and may eventually cross the blood-brain barrier, where it may infiltrate the central nervous system and causes a neuroinflammatory response. This elevated neuroinflammatory response accelerates the normal aging process in the brain, leading to neuronal damage that may make it more difficult for survivors to engage in cognitively demanding tasks. Thus, in the context of survivorship, typical processes in the aging brain are exacerbated by the accumulated cellular insults of cancer treatment, eventually leading to neurocognitive impairment later in life.

## 7. ACKNOWLEDGEMENTS

Supported by the National Cancer Institute (CA215405 [MPIs: Krull/Mandrell], CA195547 [MPIs: Hudson/Ness]; CA021765 [PI: Roberts]; CA23944 [PI: Gronemeyer]; CA239630 [MPIs: Brinkman/Krull]), and the American Lebanese-Syrian Associated Charities (ALSAC). The content is solely the responsibility of the authors and does not necessarily represent the official views of the National Institute of Health. The funding organizations had no role in the design and conduct of the study; collection, management, analysis, and interpretation of the data; preparation, review, or approval of the manuscript; and decision to submit the manuscript for publication. We thank the patients and their families for their participation.

## REFERENCES

1. Surveillance, Epidemiology, and End Results Program. SEER. Accessed July 9, 2024. https://seer.cancer.gov/index.html

2. National Cancer Institute. NCCR*Explorer: An interactive website for NCCR cancer statistics. Published online September 26, 2024. https://nccrexplorer.ccdi.cancer.gov/

3. Phillips SM, Padgett LS, Leisenring WM, et al. Survivors of Childhood Cancer in the United States: Prevalence and Burden of Morbidity. Cancer Epidemiol Biomarkers Prev. 2015;24(4):653–663. doi:10.1158/1055-9965.EPI-14-1418

4. Erdmann F, Frederiksen LE, Bonaventure A, et al. Childhood cancer: Survival, treatment modalities, late effects and improvements over time. Cancer Epidemiol. 2021;71:101733. doi:10.1016/j.canep.2020.101733

5. Bhakta N, Liu Q, Yeo F, et al. Cumulative burden of cardiovascular morbidity in paediatric, adolescent, and young adult survivors of Hodgkin’s lymphoma: an analysis from the St Jude Lifetime Cohort Study. Lancet Oncol. 2016;17(9):1325–1334. doi:10.1016/S1470-2045(16)30215-7

6. Dixon SB, Howell CR, Lu L, et al. Cardiac biomarkers and association with subsequent cardiomyopathy and mortality among adult survivors of childhood cancer: A report from the St. Jude Lifetime Cohort. Published online 2021.

7. Chow EJ, Friedman DL, Stovall M, et al. Risk of thyroid dysfunction and subsequent thyroid cancer among survivors of acute lymphoblastic leukemia: A report from the Childhood Cancer Survivor Study. Pediatr Blood Cancer. 2009;53(3):432–437. doi:10.1002/pbc.22082

8. Sklar C, Whitton J, Mertens A, et al. Abnormalities of the Thyroid in Survivors of Hodgkin’s Disease: Data from the Childhood Cancer Survivor Study. 85(9).

9. Krull KR, Cheung YT, Liu W, et al. Chemotherapy Pharmacodynamics and Neuroimaging and Neurocognitive Outcomes in Long-Term Survivors of Childhood Acute Lymphoblastic Leukemia. J Clin Oncol. 2016;34(22):2644–2653. doi:10.1200/JCO.2015.65.4574

10. Hardy KK, Hudson MM, Krull KR. Life-Altering Consequences of Neurocognitive Impairment in Survivors of Pediatric Cancer. J Clin Oncol. 2021;39(16):1693–1695. doi:10.1200/JCO.21.00211

11. Phillips NS, Baedke JL, Williams A, et al. Accelerated brain age and associated neurocognitive impairments in adult survivors of childhood cancer. J Clin Oncol. 2023;41(16_suppl):10028–10028. doi:10.1200/JCO.2023.41.16_suppl.10028

12. Phillips NS, Mulrooney DA, Williams AM, et al. Neurocognitive impairment associated with chronic morbidity in long-term survivors of Hodgkin Lymphoma. Blood Adv. 2023;7(23):7270–7278. doi:10.1182/bloodadvances.2023010567

13. Daniel S, Nylander V, Ingerslev LR, et al. T cell epigenetic remodeling and accelerated epigenetic aging are linked to long-term immune alterations in childhood cancer survivors. Clin Epigenetics. 2018;10(1):138. doi:10.1186/s13148-018-0561-5

14. Qin N, Li Z, Song N, et al. Epigenetic Age Acceleration and Chronic Health Conditions Among Adult Survivors of Childhood Cancer. JNCI J Natl Cancer Inst. 2021;113(5):597–605. doi:10.1093/jnci/djaa147

15. Ariffin H, Azanan MS, Abd Ghafar SS, et al. Young adult survivors of childhood acute lymphoblastic leukemia show evidence of chronic inflammation and cellular aging: Inflammaging in Childhood CA Survivors. Cancer. 2017;123(21):4207–4214. doi:10.1002/cncr.30857

16. Cheung YT, Brinkman TM, Mulrooney DA, et al. Impact of sleep, fatigue, and systemic inflammation on neurocognitive and behavioral outcomes in long-term survivors of childhood acute lymphoblastic leukemia: Sleep and Cognition in Childhood ALL. Cancer. 2017;123(17):3410–3419. doi:10.1002/cncr.30742

17. Krull KR, Sabin ND, Reddick WE, et al. Neurocognitive Function and CNS Integrity in Adult Survivors of Childhood Hodgkin Lymphoma. J Clin Oncol. 2012;30(29):3618–3624. doi:10.1200/JCO.2012.42.6841

18. Cheung YT, Sabin ND, Reddick WE, et al. Leukoencephalopathy and long-term neurobehavioural, neurocognitive, and brain imaging outcomes in survivors of childhood acute lymphoblastic leukaemia treated with chemotherapy: a longitudinal analysis. Lancet Haematol. 2016;3(10):e456–e466. doi:10.1016/S2352-3026(16)30110-7

19. Kesler SR, Ogg R, Reddick WE, et al. Brain Network Connectivity and Executive Function in Long-Term Survivors of Childhood Acute Lymphoblastic Leukemia. Brain Connect. 2018;8(6):333–342. doi:10.1089/brain.2017.0574

20. Kesler SR, Gugel M, Pritchard-Berman M, et al. Altered resting state functional connectivity in young survivors of acute lymphoblastic leukemia: Intrinsic Connectivity in ALL Survivors. Pediatr Blood Cancer. 2014;61(7):1295–1299. doi:10.1002/pbc.25022

21. Guida JL, Hyun G, Belsky DW, et al. Associations of seven measures of biological age acceleration with frailty and all-cause mortality among adult survivors of childhood cancer in the St. Jude Lifetime Cohort. Nat Cancer. 2024;5(5):731–741. doi:10.1038/s43018-024-00745-w

22. Williams AM, Mandelblatt J, Wang M, et al. Premature aging as an accumulation of deficits in young adult survivors of pediatric cancer. JNCI J Natl Cancer Inst. 2023;115(2):200–207. doi:10.1093/jnci/djac209

23. Williams AM, Mandelblatt JS, Wang M, et al. Deficit Accumulation Index and Biological Markers of Aging in Survivors of Childhood Cancer. JAMA Netw Open. 2023;6(11):e2344015. doi:10.1001/jamanetworkopen.2023.44015

24. Ding XQ, Maudsley AA, Sabati M, et al. Physiological neuronal decline in healthy aging human brain — An in vivo study with MRI and short echo-time whole-brain 1H MR spectroscopic imaging. NeuroImage. 2016;137:45–51. doi:10.1016/j.neuroimage.2016.05.014

25. Boumezbeur F, Mason GF, De Graaf RA, et al. Altered Brain Mitochondrial Metabolism in Healthy Aging as Assessed by *in vivo* Magnetic Resonance Spectroscopy. J Cereb Blood Flow Metab. 2010;30(1):211–221. doi:10.1038/jcbfm.2009.197

26. Kantarci K, Weigand SD, Petersen RC, et al. Longitudinal 1H MRS changes in mild cognitive impairment and Alzheimer’s disease. Neurobiol Aging. 2007;28(9):1330–1339. doi:10.1016/j.neurobiolaging.2006.06.018

27. Haga KK, Khor YP, Farrall A, Wardlaw JM. A systematic review of brain metabolite changes, measured with 1H magnetic resonance spectroscopy, in healthy aging. Neurobiol Aging. 2009;30(3):353–363. doi:10.1016/j.neurobiolaging.2007.07.005

28. Howell CR, Bjornard KL, Ness KK, et al. Cohort Profile: The St. Jude Lifetime Cohort Study (SJLIFE) for paediatric cancer survivors. Int J Epidemiol. 2021;50(1):39–49. doi:10.1093/ije/dyaa203

29. Bottomley PA. Spatial Localization in NMR Spectroscopy in Vivo. Ann N Y Acad Sci. 1987;508(1):333–348. doi:10.1111/j.1749-6632.1987.tb32915.x

30. Di X, Rypma B, Biswal BB. Correspondence of executive function related functional and anatomical alterations in aging brain. Prog Neuropsychopharmacol Biol Psychiatry. 2014;48:41–50. doi:10.1016/j.pnpbp.2013.09.001

31. Brosnan MB, Wiegand I. The Dorsolateral Prefrontal Cortex, a Dynamic Cortical Area to Enhance Top-Down Attentional Control. J Neurosci. 2017;37(13):3445–3446. doi:10.1523/JNEUROSCI.0136-17.2017

32. Raz N, Rodrigue KM, Haacke EM. Brain Aging and Its Modifiers: Insights from *in Vivo* Neuromorphometry and Susceptibility Weighted Imaging. Ann N Y Acad Sci. 2007;1097(1):84–93. doi:10.1196/annals.1379.018

33. Duff MC, Brown-Schmidt S. The hippocampus and the flexible use and processing of language. Front Hum Neurosci. 2012;6. doi:10.3389/fnhum.2012.00069

34. Yassa MA, Reagh ZM. Competitive Trace Theory: A Role for the Hippocampus in Contextual Interference during Retrieval. Front Behav Neurosci. 2013;7. doi:10.3389/fnbeh.2013.00107

35. Bone MB, Buchsbaum BR. Concurrent Feature-Specific Reactivation within the Hippocampus and Neocortex Facilitates Episodic Memory Retrieval. Neuroscience; 2021. doi:10.1101/2021.02.28.433226

36. Duff MC, Covington NV, Hilverman C, Cohen NJ. Semantic Memory and the Hippocampus: Revisiting, Reaffirming, and Extending the Reach of Their Critical Relationship. Front Hum Neurosci. 2020;13:471. doi:10.3389/fnhum.2019.00471

37. Stoodley CJ, Valera EM, Schmahmann JD. Functional topography of the cerebellum for motor and cognitive tasks: An fMRI study. NeuroImage. 2012;59(2):1560–1570. doi:10.1016/j.neuroimage.2011.08.065

38. Burman R, Guthrie S, Pang Y, et al. Building a standalone automated High-Throughput LCModel prototype application for 1H-MRS data processing: HT-LCModel. Published online 2025.

39. Provencher SW. Automatic quantitation of localized in vivo 1H spectra with LCModel. NMR Biomed. 2001;14(4):260–264. doi:10.1002/nbm.698

40. Delis DC, Kramer JH, Kaplan E. Integrating Clinical Assessment With Cognitive Neuroscience: Construct Validation of the California Verbal Learning Test.

41. Hilbert S, Nakagawa TT, Puci P, Zech A, Bühner M. The Digit Span Backwards Task: Verbal and Visual Cognitive Strategies in Working Memory Assessment. Eur J Psychol Assess. 2015;31(3):174–180. doi:10.1027/1015-5759/a000223

42. Joy S. Speed and memory in the WAIS-III Digit Symbol?Coding subtest across the adult lifespan. Arch Clin Neuropsychol. 2004;19(6):759–767. doi:10.1016/j.acn.2003.09.009

43. Tombaugh T. Trail Making Test A and B: Normative data stratified by age and education. Arch Clin Neuropsychol. 2004;19(2):203–214. doi:10.1016/S0887-6177(03)00039-8

44. Tombaugh TN, Kozak J, Rees L. Normative Data Stratified by Age and Education for Two Measures of Verbal Fluency: FAS and Animal Naming. Published online 1999.

45. R Core Team. R: A language and environment for statistical computing. R Found Stat Comput. Published online 2018:14.

46. Mahmoudi N, Dadak M, Bronzlik P, et al. Microstructural and Metabolic Changes in Normal Aging Human Brain Studied with Combined Whole-Brain MR Spectroscopic Imaging and Quantitative MR Imaging. Clin Neuroradiol. Published online June 19, 2023. doi:10.1007/s00062-023-01300-3

47. Maghsudi H, Schütze M, Maudsley AA, Dadak M, Lanfermann H, Ding XQ. Age-related Brain Metabolic Changes up to Seventh Decade in Healthy Humans: Whole-brain Magnetic Resonance Spectroscopic Imaging Study. Clin Neuroradiol. 2020;30(3):581–589. doi:10.1007/s00062-019-00814-z

48. Moscovitch M, Cabeza R, Winocur G, Nadel L. Episodic Memory and Beyond: The Hippocampus and Neocortex in Transformation. Annu Rev Psychol. 2016;67(1):105–134. doi:10.1146/annurev-psych-113011-143733

49. Preston AR, Eichenbaum H. Interplay of Hippocampus and Prefrontal Cortex in Memory. Curr Biol. 2013;23(17):R764–R773. doi:10.1016/j.cub.2013.05.041

50. Song T, Song X, Zhu C, et al. Mitochondrial dysfunction, oxidative stress, neuroinflammation, and metabolic alterations in the progression of Alzheimer’s disease: A meta-analysis of in vivo magnetic resonance spectroscopy studies. Ageing Res Rev. 2021;72:101503. doi:10.1016/j.arr.2021.101503

51. Williams AM, Liu W, Ehrhardt MJ, et al. Systemic Biological Mechanisms of Neurocognitive Dysfunction in Long-Term Survivors of Childhood Hodgkin Lymphoma. Clin Cancer Res. 2024;30(9):1822–1832. doi:10.1158/1078-0432.CCR-23-3709

52. Brand A, Richter-Landsberg C, Leibfritz D. Multinuclear NMR studies on the energy metabolism of glial and neuronal cells. Dev Neurosci. 1993;15(3-5):289–298. doi:10.1159/000111347

53. Feldman RA. Microglia orchestrate neuroinflammation. eLife. 2022;11:e81890. doi:10.7554/eLife.81890

54. Hyder F, Patel AB, Gjedde A, Rothman DL, Behar KL, Shulman RG. Neuronal–Glial Glucose Oxidation and Glutamatergic–GABAergic Function. J Cereb Blood Flow Metab. 2006;26(7):865–877. doi:10.1038/sj.jcbfm.9600263

55. Rothman DL, Behar KL, Hyder F, Shulman RG. In vivo NMR Studies of the Glutamate Neurotransmitter Flux and Neuroenergetics: Implications for Brain Function. Annu Rev Physiol. 2003;65(1):401–427. doi:10.1146/annurev.physiol.65.092101.142131

56. Williams AM, Mirzaei Salehabadi S, Xing M, et al. Modifiable risk factors for neurocognitive and psychosocial problems after Hodgkin lymphoma. Blood. 2022;139(20):3073–3086. doi:10.1182/blood.2021013167

57. Cheung YT, Khan RB, Liu W, et al. Association of Cerebrospinal Fluid Biomarkers of Central Nervous System Injury With Neurocognitive and Brain Imaging Outcomes in Children Receiving Chemotherapy for Acute Lymphoblastic Leukemia. JAMA Oncol. 2018;4(7):e180089. doi:10.1001/jamaoncol.2018.0089

58. Ikonomidou C. Chemotherapy and the pediatric brain. Mol Cell Pediatr. 2018;5(1):8. doi:10.1186/s40348-018-0087-0

59. Mescher M, Merkle H, Kirsch J, Garwood M, Gruetter R. Simultaneousin vivo spectral editing and water suppression. NMR Biomed. 1998;11(6):266–272. doi:10.1002/(SICI)1099-1492(199810)11:6%3C266::AID-NBM530%3E3.0.CO;2-J

60. Janssen B, Vugts D, Windhorst A, Mach R. PET Imaging of Microglial Activation—Beyond Targeting TSPO. Molecules. 2018;23(3):607. doi:10.3390/molecules23030607

61. Largeau B, Dupont AC, Guilloteau D, Santiago-Ribeiro MJ, Arlicot N. TSPO PET Imaging: From Microglial Activation to Peripheral Sterile Inflammatory Diseases? Contrast Media Mol Imaging. 2017;2017:1–17. doi:10.1155/2017/6592139

62. Zhang L, Hu K, Shao T, et al. Recent developments on PET radiotracers for TSPO and their applications in neuroimaging. Acta Pharm Sin B. 2021;11(2):373–393. doi:10.1016/j.apsb.2020.08.006

63. Qiao L, Fisher E, McMurray L, et al. Radiosynthesis of (*R*, *S*)-[ ^18^ F]GE387: A Potential PET Radiotracer for Imaging Translocator Protein 18 kDa (TSPO) with Low Binding Sensitivity to the Human Gene Polymorphism rs6971. ChemMedChem. 2019;14(9):982–993. doi:10.1002/cmdc.201900023

